# Extending PROXIMAL to predict degradation pathways of phenolic compounds in the human gut microbiota

**DOI:** 10.1101/2023.05.17.541107

**Authors:** Francesco Balzerani, Telmo Blasco, Sergio Pérez-Burillo, Luis Valcarcel, Francisco J. Planes, Soha Hassoun

## Abstract

Despite significant advances in reconstructing genome-scale metabolic networks, the understanding of cellular metabolism remains incomplete for many organisms. A promising approach for elucidating cellular metabolism is analysing the full scope of enzyme promiscuity, which exploits the capacity of enzymes to bind to non-annotated substrates and generate novel reactions. To guide time-consuming costly experimentation, different computational methods have been proposed for exploring enzyme promiscuity. One relevant algorithm is PROXIMAL, which strongly relies on KEGG to define generic reaction rules and link specific molecular substructures with associated chemical transformations. Here, we present a completely new pipeline, PROXIMAL2, which overcomes the dependency on KEGG data. In addition, PROXIMAL2 introduces two relevant improvements with respect to the former version: i) correct treatment of multi-step reactions and ii) tracking of electric charges in the transformations. We compare PROXIMAL and PROXIMAL2 in recovering annotated products from substrates in KEGG reactions, finding a highly significant improvement in the level of accuracy. We then applied PROXIMAL2 to predict degradation reactions of phenolic compounds in the human gut microbiota. The results were compared to RetroPath RL, a different and relevant enzyme promiscuity method. We found a significant overlap between these two methods but also complementary results, which open new research directions into this relevant question in nutrition.

## Introduction

Metabolism is defined as the whole set of chemical reactions that take place in organisms (Blanco and Blanco, 2017). In particular, metabolic pathways represent chemical transformations where a substrate becomes a product, typically with the aid of other molecules such as cofactors (Hafner and Hatzimanikatis, 2021; Folador *et al*., 2019). Recent advances in sequencing technologies have significantly increased the coverage of metabolic pathways in dozens of organisms. Much effort has been done to integrate these metabolic pathway into genome-scale metabolic models (GEMMs) (Thiele *et al*., 2013), which aim to accurately define the stoichiometry of all reactions in a particular organism, their associated genes, enzymes or transporters, their compartment localization and other relevant biological information. GEMMs allow us to analyse the metabolic capabilities of both unicellular and multicellular systems with computational tools developed in the field of constraint-based modelling (Price *et al*., 2003). Despite these advances, the understanding of cellular metabolism in many organisms is still incomplete, with significant gaps and metabolites that have no links to any reaction in available metabolic models (MohammadiPeyhani *et al*., 2022; Amin *et al*., 2019). For example, even in the well-annotated KEGG database (Kanehisa and Goto, 2000), there remains 10,000 metabolites that are still not linked to a known biochemical reaction (MohammadiPeyhani *et al*., 2022).

Different computational tools have been developed to fill in metabolic gaps by means of improving the functional annotation of enzymes. The number of annotated enzymes across databases is much lower than that of metabolic reactions, which could be pointing towards enzymes carrying out more than one biochemical reaction. In KEGG, while we have 11,822 reactions, we can only find 8,012 enzymes (https://www.kegg.jp/kegg/docs/statistics.html). The same pattern is found in BRENDA (Jeske *et al*., 2019), where the numbers are 21,665 and 8,332, respectively. Similarly, in AGORA (Magnúsdóttir *et al*., 2017), a repository of metabolic networks that include 818 organisms from the human gut microbiota, we found a total count of 1438.1 reactions on average per organism. In the case of enzymes, considering the genomic annotation from GenBank (Benson *et al*., 2017) and Ensembl (Kersey *et al*., 2018), we obtained a total of 900.7 enzymes on average per organism in AGORA. Considering this evidence, annotating enzyme promiscuity seems a promising strategy to improve metabolic networks.

Promiscuous activity of enzymes lies on their capacity to bind to non-canonical substrates and catalyse novel reactions (Gupta, 2016; Copley, 2017). Annotating such capabilities provides the possibility to understand underground metabolism, which is not represented in current databases (Notebaart *et al*., 2018; Guzmán *et al*., 2019). The growing relevance on enzyme promiscuity in different fields has led to the development of a number of algorithms and computational methods (Carbonell, Parutto, Herisson, *et al*., 2014; Carbonell and Faulon, 2010; Kumar *et al*., 2018; MohammadiPeyhani *et al*., 2022). In brief, these algorithms rely on a set of generic enzymatic reaction rules, which define the chemical transformations that occur to a substrate, describing its reactive site and atomic rearrangement as a result of the reaction (Koch *et al*., 2020; MohammadiPeyhani *et al*., 2022). These reactions rules describe an abstraction of known reactions and permit a certain degree of flexibility of the substrates involved (Ni *et al*., 2021), potentially leading to new reactions and products.

Several methods use manually curated reaction rules (Li *et al*., 2004; Hadadi *et al*., 2016; Hafner *et al*., 2020; Jeffryes *et al*., 2015). Although these rules integrate the best knowledge about enzymes, they are limited to a reduced number of reactions (Ni *et al*., 2021). For this reason, the development of computational tools able to automatically extract reaction rules from known transformations has received much attention. Considerable progress has been made in recent years (Duigou *et al*., 2019; Yousofshahi *et al*., 2015; Kumar *et al*., 2018; Ni *et al*., 2021). RetroRules (Duigou *et al*., 2019) provides thousands of rules that are extracted from public databases and constitutes the core of different algorithms for predicting novel metabolic pathways, such as RetroPath RL (Koch *et al*., 2020), the latest version of a series of works developed by the same authors (Carbonell, Parutto, Baudier, *et al*., 2014; Delépine *et al*., 2018; Duigou *et al*., 2019; Koch *et al*., 2020).

In a previous effort to elucidate the metabolism of phenolic compounds in the human gut microbiota, we applied RetroPath RL to predict phenolic degradation pathways in the human gut microbiota. As more than 2/3 of phenolic compounds in the Phenol Explorer database (Rothwell *et al*., 2013) are not included in universal metabolic databases, it was necessary to employ computational approaches to uncover microbial phenol metabolism. Despite identifying degradation pathways for 80 compounds in the Phenol Explorer database that were not present in previous gut microbiota reconstructions, RetroPath RL could not find candidate pathways for 180 out of 372 of phenolic compounds in the Phenol Explorer database. Continuing this early work, we focus here on PROXIMAL (Yousofshahi *et al*., 2015), an enzyme promiscuity algorithm that follows a different strategy to build reaction rules and, thus, could potentially complement the results obtained with RetroPath RL.

The PROXIMAL algorithm has been successfully applied to create extended metabolic models in different organisms and to annotate cellular products (Amin *et al*., 2019; Hassanpour *et al*., 2020). PROXIMAL makes use of the KEGG database to predict possible transformations (Kanehisa and Goto, 2000). In particular, it is based on RPAIRS (Kotera *et al*., 2004), a database available in KEGG that provides the necessary alignment between paired substrates and products to define the modified sub-structures. The main limitation of this algorithm is that it cannot be used for transformations not included in KEGG. Moreover, RPAIRS was discontinued in 2016 (https://www.genome.jp/kegg/kegg1a.html), which hampers the application of PROXIMAL to more recent updated versions of KEGG, that is in continuous development. Overall, these limitations restrict the application of PROXIMAL to our problem of phenolic compound degradation in the human gut microbiota, since we rely on AGREDA (Blasco *et al*., 2021), a metabolic reconstruction that contains relevant reactions not included in KEGG.

To address these issues, we present here a completely new pipeline, called PROXIMAL2, which overcomes the dependency on KEGG data. Moreover, PROXIMAL2 extends the previous methodology for the automatic reaction rule generation, which was unable to correctly model complex reactions involved in the phenolic compound metabolism. In particular, PROXIMAL2 introduces two relevant improvements with respect to the former version: i) correct treatment of multi-step reactions and ii) tracking of electric charges in the transformations. We show that PROXIMAL2 substantially extends the chemical space of PROXIMAL, and it correctly generates a higher number of reaction rules in KEGG. We also present the application of PROXIMAL2 to predict degradation pathways of phenolic compounds in the human gut microbiota and compare the results with RetroPath RL.

## Methods

PROXIMAL (Yousofshahi *et al*., 2015) is a rule-based method for the prediction of metabolic products. PROXIMAL defines look-up tables for a database of substrate-product pairs. These look-up tables link specific molecular substructures with their associated chemical transformations (Figure 1a). In particular, ‘keys’ in the look-up tables specify the modified substrate structure, including: i) the reaction centre, atom where the chemical transformation occurs; ii) adjacent neighbours, atoms connected to the reaction centre at distance 1; iii) distant neighbours, atoms connected to the reaction centre at distance 2. In other words, the modified substrate structure induces a subgraph of neighbours within radius 2 starting from the reaction centre. In addition, ‘values’ in the look-up tables describe the modifications resulting in the product, including: i) reaction centre, ii) adjacent neighbours and iii) added/removed functional group, which defines the modified part of the substrate. Note here that the atoms in the look-up tables follow the nomenclature of ‘KEGG atom types’, which are defined according to their functional groups and microenvironment, *e.g.* C8x or C8y.

**Figure 1:**
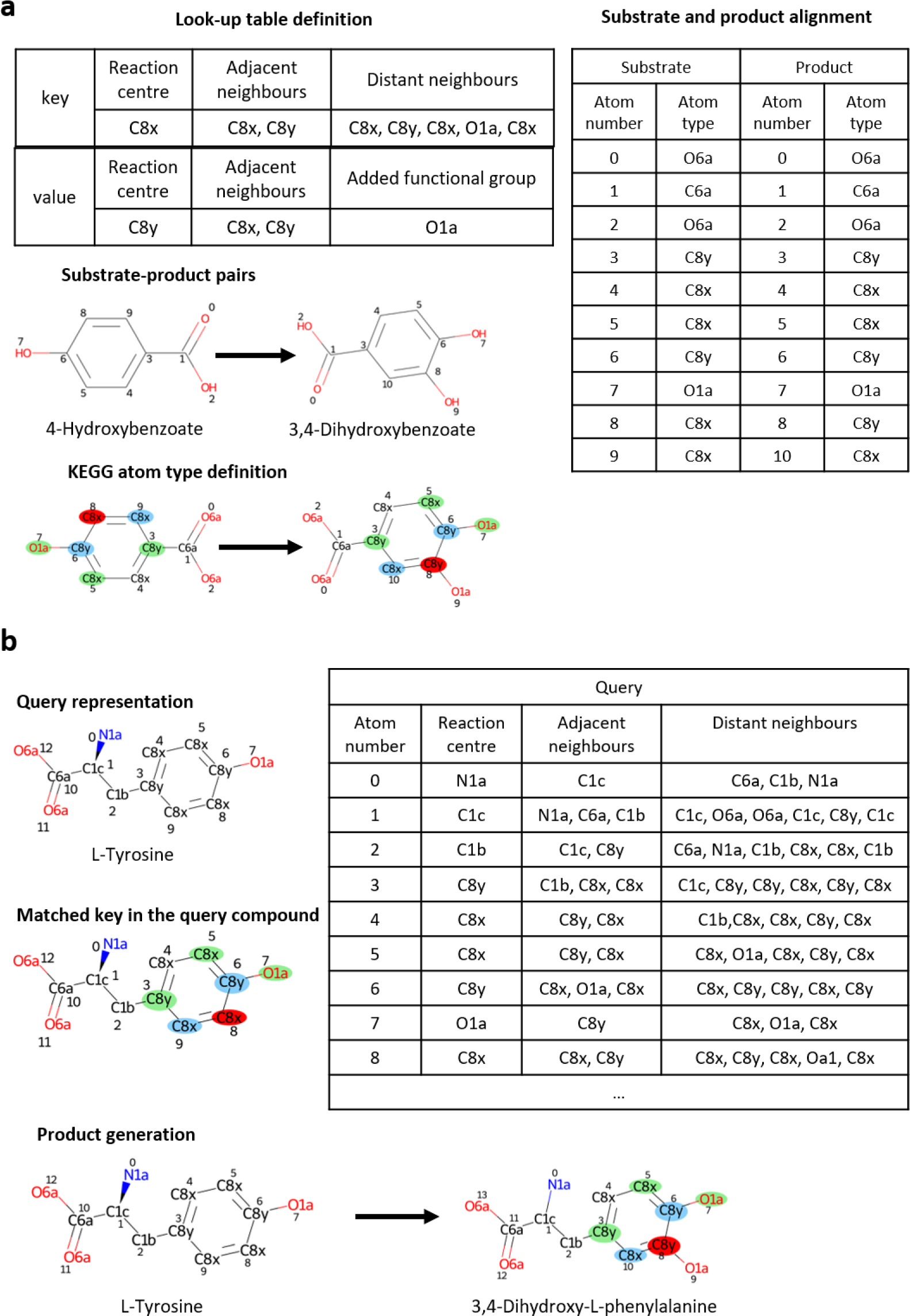
Schematic representation of the PROXIMAL workflow. **a)** Definition of an example look-up table in PROXIMAL. After the pair selection, the substrate and product are represented in KEGG atom type and then aligned to permit the definition of the reaction centre and its neighbours up to distance 2. The starting reaction is (*R01296)*: *4-Hydroxybenzoate + O2 + NADH + H(+)-> 3,4-Dihydroxybenzoate + H2O + NAD(+).* **b)** Application of the look-up table to an example selected query. The latter is represented in KEGG atom type and the key of the look-up table is searched within the molecule. Finally, the product is generated based on the look-up table value. The red, blue and green circles represent the reaction centre, adjacent and distant neighbour, respectively. The atoms O and N are marked in red and blue, respectively, as conventionally used in chemistry by the CPK colouring rules.

Once these look-up tables are defined for all substrate-product pairs involved in the database of reactions of interest, PROXIMAL lists the different subgraphs of neighbours within radius 2 for a given query compound, searches for the ones matching with the sub-structures stored in the key tables and applies their associated transformation defined in the value tables in order to generate putative products. Figure 1b illustrates that the subgraph of neighbours centred at atom number 8 in the query compound L-Tyrosine matches with the key table defined in Figure 1a for the pair 4-Hydroxybenzoate and 3,4-Dihydroxybenzoate, leading to the compound 3,4-Dihydroxy-L-phenylalanine.

The identification of the reaction centre for each substrate-product pair is the critical step in the definition of look-up tables mentioned above. This is done following different steps that require i) the alignment of the substrate and product atoms and ii) the classification of different atoms into different ‘KEGG atom types’ (Figure 1a). Reaction centres are defined as any specific substrate atom that aligns with a product atom of different KEGG atom type. In Figure 1a, the reaction centre corresponds with the atom number 8 of 4-Hydroxybenzoate and the atom number 8 of 3,4-Dihydroxybenzoate, whose KEGG atom types are C8x and C8y, respectively.

The look-up tables in PROXIMAL were built using the RPAIRS database, where substrate-product pairs are defined for each reaction, atomic species are classified into KEGG atom types, and substrate atoms are aligned to product atoms. Unfortunately, as noted above, RPAIRS database was discontinued in 2016, which restricts its application to a limited subset of reactions and, thus, metabolic space (*i.e.* set of potential biochemical transformations and molecules (Carbonell, Parutto, Herisson, *et al*., 2014)). Therefore, we present here a completely new pipeline, called PROXIMAL2, summarized in Figure 2a.

**Figure 2:**
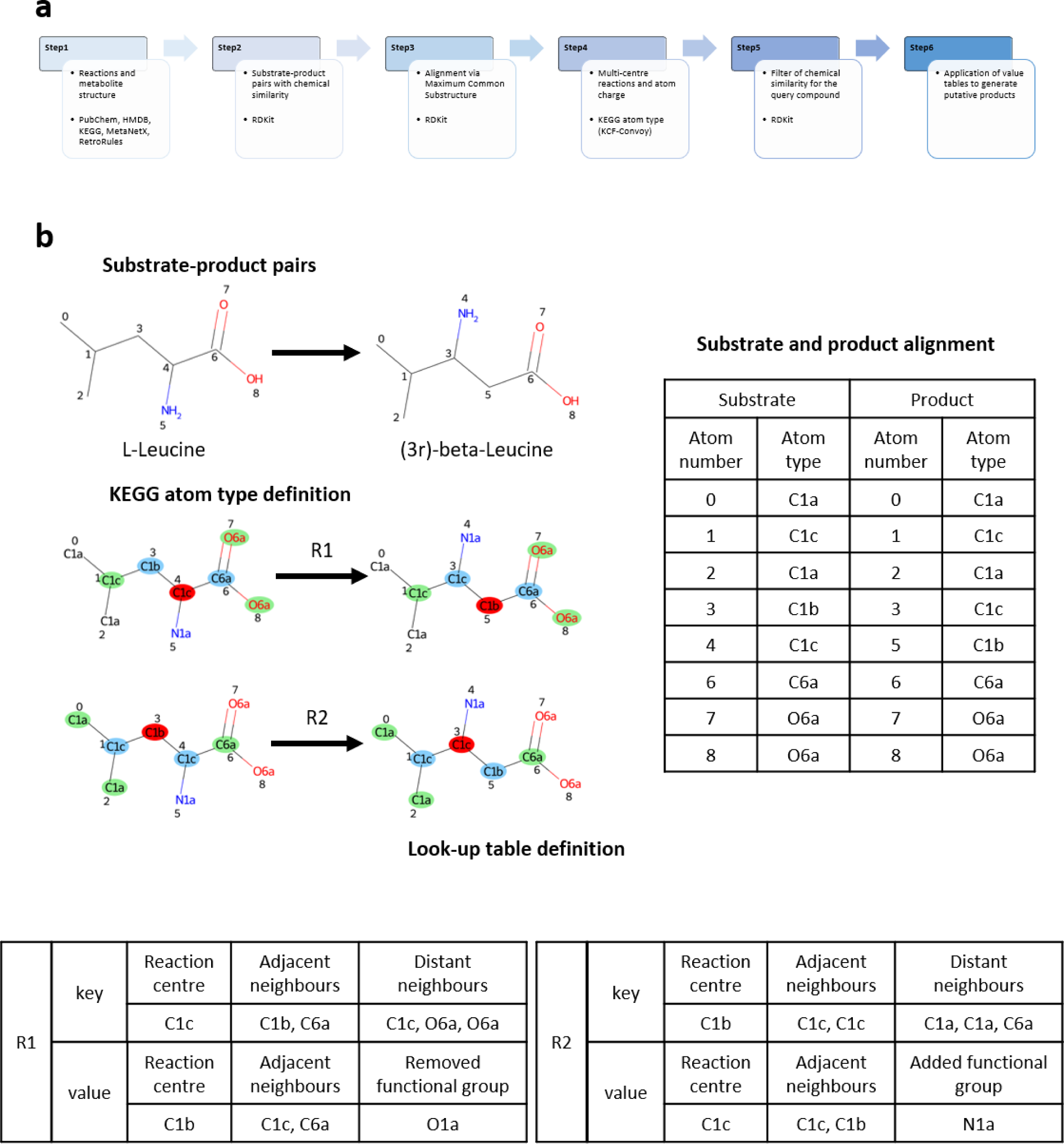
PROXIMAL2 workflow and illustration of multi-centre reactions. **a)** PROXIMAL2 pipeline summarizing main changes with respect to the previous version. **b)** Example of multi-centre reactions. The alignment provides two changing KEGG atom types (atom numbers 3 and 4), leading to two reaction centre that are labelled as *R1* and *R2*. In the look-up table, the information about the reaction centres and respective neighbour is stored. The reaction is (*R01091)*: *L-Leucine-> (3R)-beta-Leucine*.

### Step 1: Definition of the database of reactions, metabolites and structural information

First, we created the metabolic space that is required to apply our enzyme promiscuity algorithm. Starting from an input database of reactions, we first removed the most common cofactors involved within the transformations and deleted the reactions if they only included cofactors. Then, the metabolites involved in the different transformations were extracted. We obtained SMILES and InChI for each metabolite from different public databases: PubChem (Kim *et al*., 2019), KEGG (Kanehisa and Goto, 2000), HMDB (Wishart *et al*., 2018), MetaNetX (Moretti *et al*., 2016) and RetroRules (Duigou *et al*., 2019). Metabolites with no available structure were filtered out from the reactions; similarly, input reactions only involving metabolites without structure were deleted. As a result, we obtained a list of simplified reactions.

### Step 2: Definition of substrate-product pairs

As noted above, PROXIMAL works with substrate-product pairs. However, the output (simplified) reactions from Step1 can include more than one substrate or product. To identify the best matching pairs for a given reaction, we calculated the chemical similarity between each substrate-product pair and paired them according to such value. The chemical similarity was determined using the RDKit package (Landrum, 2011) and the Morgan Fingerprint (Rogers and Hahn, 2010). In reactions with a single substrate (or product), the latter is associated to each product (substrate) to form more than one pair. The output of this step is the list of substrate-product pairs for the generation of look-up tables. Note that with this chemoinformatic approach we recovered 97.5% of pairs annotated in RPAIRS.

### Step 3: Alignment of substrate-product pairs

As noted above, the identification of the reaction centre and look-up tables for each substrate-product pair requires their atomic alignment. To that end, we extracted the maximum common substructure (MCS) between the substrate and product with the function *findMCS* available in the module *rdFMCS* in RDKit package. We fixed two optional parameters in this RDKit function in order to consider the specificity of the atoms belonging to rings and charged atoms. In particular, a match between atoms included in rings can happen only if the atoms are part of rings in both the substrate and product. Similarly, two atoms can match with each other only when both have the same charge (Supplementary Methods, Supplementary Figure 1).

### Step 4: Definition of reaction centres and look-up tables

As noted above, reaction centres are defined as any specific substrate atom that aligns with a product atom of different KEGG atom type. Here, the classification of substrate and product atoms into different KEGG atom types were done with the package KCF-Convoy (Kotera *et al*., 2013; Sato *et al*., 2018). With this information and the atomic alignment (Step3), reaction centres were identified.

A relevant improvement included in PROXIMAL2 is the possibility to analyse reactions containing multiple reaction centres. In some cases, it may happen that a reaction represents a multi-step reaction (Zhou and Zhuang, 2007), where the intermediate steps are removed for any reason (*e.g.* inability to measure the intermediate), and they are unified in the transformation that connects the initial substrates directly to the final products. This concept translates into having more than one reaction centre when the changing bonds are analysed. For each reaction centre involved in these transformations, look-up tables were defined, as shown in Figure 2b.

In addition, in some cases, we found a modification of the bonds between the atoms within the MCS, showing a new arrangement due to the transformation (Supplementary Methods). Specifically, during the rearrangement of the structure, a bond can be introduced, deleted or simply changed (*e.g.* going from a double to single bond). These modifications were also extracted together and included in the look-up tables as multiple reaction centres (Supplementary Figure 2). Finally, the information about the possible position of a charged atom is extracted in order to apply that charge when the transformation is applied.

### Step 5: Search of matching keys within the query compound

Once the look-up tables representing the chemical transformations are generated, we define the subgraph of neighbours within radius 2 for each atom of the query compound and search for matching keys in the look-up tables. For multi-centre reactions we ensure that the query compound matches with all their associated keys. Note here that a pre-filter was implemented to any operator applied to the query compound. In particular, we discard matches where the chemical similarity between the query compound and the substrate of the matching entry was below 0.6.

### Step 6: Product generation

Once an operator in the look-up table matches the query compound, we generated the promiscuous product. Therefore, considering the molecule of interest, the transformation is applied to each reaction centre, adding atoms and bonds coherently according to the original template reaction. When there is more than one reaction centre, it can happen that a substructure, which must be added to the query, is in common between several reaction centres. To avoid the addition of that substructure multiple times, we implemented a tracking system of the atoms and bonds already introduced, rejecting duplicate additions. Then, bonds are added, removed or changed coherently to the operator definition. Finally, information about the charges present in the query is kept and introduced in the generated product molecule. If a charged atom was removed due to the transformation, the charge will not be present in the final product. Similarly, when a functional group introduced in the structure contained a charged atom, that charge was introduced in the predicted molecule.

Once all the changes within the chemical transformation are applied, the generated products are saved as json file, where the information about the final product (Smiles ID and mol block text), the reaction template (ID, EC number and Formula), and the initial molecule (name, ID and structure) is stored. Although the generation of the mol block text was always possible because it was generated manually, the SMILES string was generated using the RDKit package.

## Results

### Comparison between PROXIMAL and PROXIMAL2

We first evaluated the accuracy of PROXIMAL2 in recovering annotated products from substrates in KEGG reactions, in comparison with the previous PROXIMAL algorithm (https://hassounlab.cs.tufts.edu/proximal/).

Therefore, we applied PROXIMAL2 to the same set of KEGG reactions used by PROXIMAL This subset of reactions involves 8819 reactions and 4983 associated metabolites (Supplementary Tables 1-2). We derived the key and value tables for all the substrate-product pairs, following the methodology presented in the Methods section, and investigated the capacity to generate the annotated product for each of these pairs when the associated substrate is used as a query compound. We compared the accuracy of PROXIMAL2 with PROXIMAL (see Figure 3a).

**Figure 3:**
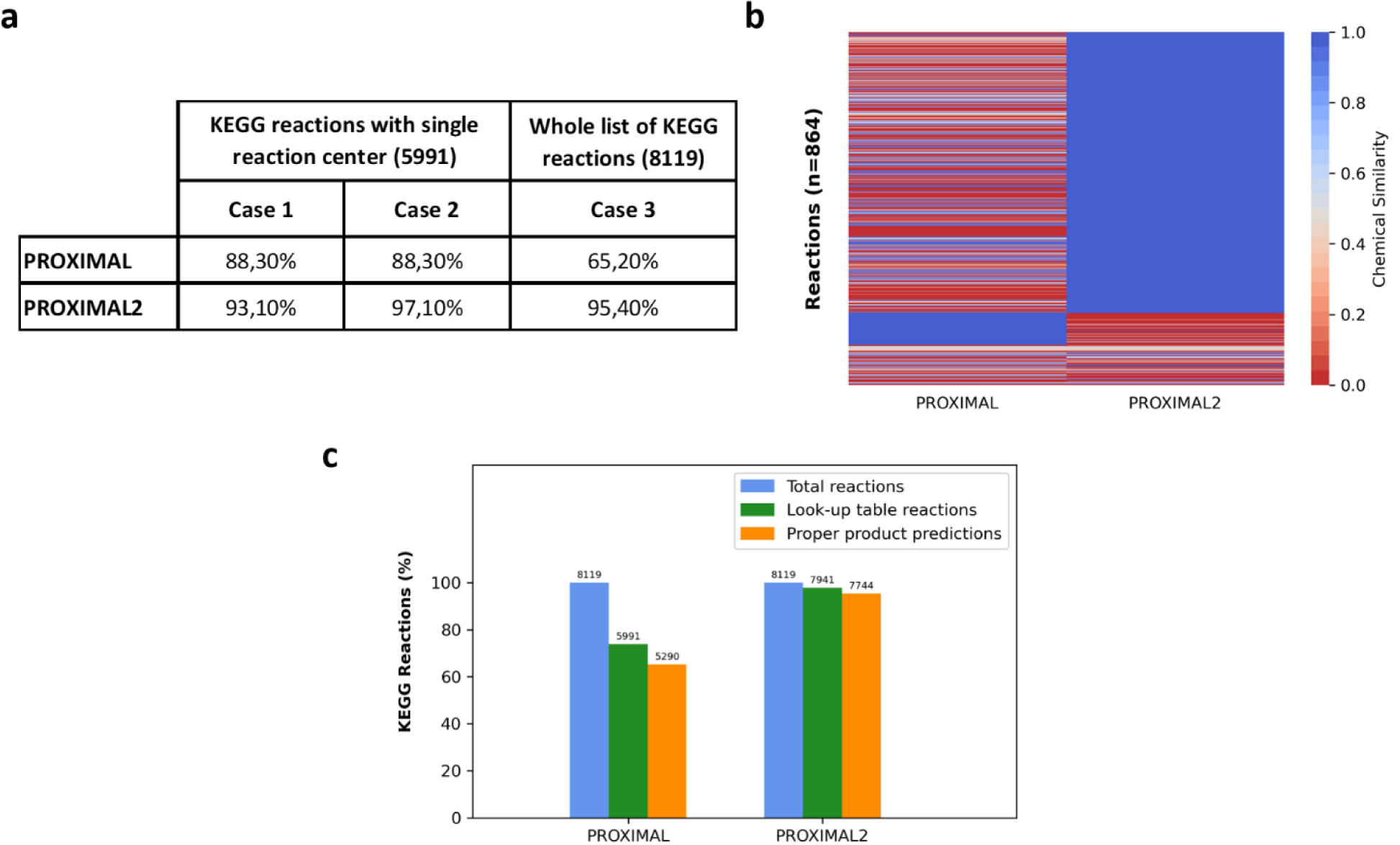
Comparison between PROXIMAL and PROXIMAL2. **a)** Percentage accuracy in recovering annotated products in KEGG for PROXIMAL and PROXIMAL2 in three different scenarios: Case1, Case2 and Case3. In Case1 and Case2, we consider reactions with a single reaction centre. We do not consider atom charge information in Case1, but we do in Case2. In Case3, we consider all the reactions in the KEGG version used in PROXIMAL. The number of reactions used in each of the cases is shown in parenthesis. **b)** Heatmap representing the chemical similarity of the predicted and annotated products in KEGG in Case 1 and 2 where at least one of the two algorithms fail to predict the annotated product. **c)** Barplot representing the reaction coverage with look-up tables and those correctly predicting the annotated product in KEGG reactions. The y-axis shows the coverage in percentage. The total number of reactions are indicated over the bars.

First, we analysed the performance of PROXIMAL2 with the same limitations as in PROXIMAL. In particular, we only considered those reactions with a single reaction centre, reducing the study to 5991 reactions, and neglected the atom charge information from the predictions of PROXIMAL2. Under this scenario (Case 1 in Figure 3a), PROXIMAL2 was able to generate the proper product for 5574 out of 5991 reactions (accuracy: 93%), while the predictions were correct for 5290 reactions in PROXIMAL (accuracy: 88.3%). This result illustrates that our chemoinformatic strategy in PROXIMAL2 (Steps 1-3), which overcomes the dependence on RPAIRS, produces more accurate results than PROXIMAL. In order to evaluate the effect of considering atom charge in PROXIMAL2, we updated the previous comparison and included this feature in our analysis. We now reached the proper product for 5814 reactions, obtaining an accuracy of 97% (Case 2 in Figure 3a). This shows that the effect of atom charge in PROXIMAL2 further increase the accuracy of PROXIMAL2.

To visualize the differences between PROXIMAL and PROXIMAL2 in Case 2, we extracted the molecules for which at least one of the two algorithms were not able to produce the correct product and computed the chemical similarity values between the predicted and annotated product (Figure 3b). It can be observed that PROXIMAL2 is substantially more accurate than PROXIMAL, finding the annotated product (chemical similarity = 1) in many more cases. Note here that in both cases, Case 1 and Case 2, the improvement of PROXIMAL2 is highly significant (two proportions z-test p-value ≤ 2e-16).

Finally, we included multi-centre reactions in our analysis and considered the whole set of 8119 reactions. PROXIMAL could not improve the accuracy, since it is not able to model multi-centre reactions, obtaining an accuracy of 65.2%. PROXIMAL2, instead, generated the correct product for 7744 reactions (accuracy: 95.4%), which illustrates the clear advance brought by PROXIMAL2 (Case 3 in Figure 3a). Note here that we could only generate look-up tables for 7941 out of 8119 reactions with PROXIMAL2 and, thus, the accuracy is even higher for these subset of reactions (97.5%). In both cases, the improvement of PROXIMAL2 is statistically significant (two proportions z-test p-value ≤ 2e-16). PROXIMAL2 was not able to generate look-up tables for the remainder 178 reactions mainly due to the incapacity to deal with stereochemical information and the restrictions of atom charge imposed in the definition of MCS (see Supplementary Methods).

### Application of PROXIMAL2 to predict the degradation of phenolic compounds in the human gut microbiota

We defined the metabolic space of the human gut microbiota, particularly relevant for the degradation of phenolic compounds ingested from diet, based on our recently published metabolic reconstruction called AGREDA (Blasco *et al*., 2021). Specifically, we started our analysis from the set of 5087 reactions used in our previous work (Balzerani *et al*., 2022) (Supplementary Table 3), where the same question was addressed with a different enzyme promiscuity algorithm, RetroPath RL.

RetroPath RL is a rule-based method that makes use of the RetroRules database (Duigou *et al*., 2019) to investigate enzyme promiscuity. RetroRules defines the reaction centre based on an atom-atom mapping between substrates and product atoms and compute reaction rules using the reaction SMARTS (SMILES Arbitrary Target Specification) formalism. The level of specificity of reaction rules in RetruRules can be tuned according to the diameter parameter, *i.e.* the size of a hypothetic sphere around the reaction centre. Moreover, RetroPath RL allows the user to fix the threshold of chemical similarity between the query compound and the substrate of the rule. To apply RetroPath RL in the most similar conditions to PROXIMAL2, we fixed the diameter at 4 and the chemical similarity threshold at 0.6.

A high proportion of reactions considered in the metabolic space of AGREDA were present in the RetroRules database; however, we also included some manually curated reactions important for the metabolism of phenolic compounds (Blasco *et al*., 2021; Balzerani *et al*., 2022). PROXIMAL2 only requires the biochemical reaction equation to generate the look-up tables (reaction rules). For the reactions present in RetroRules, we extracted the equations from MetaNetX database, whereas the equations for the manually curated reactions were obtained from AGREDA. Following the complete pipeline of PROXIMAL2, we could generate look-up tables (reaction rules) for 4860 reactions (Figure 4a). Reaction rules for RetroPath RL, in contrast, were directly obtained from the RetroRules database at diameter 4. For manually curated reactions, we had to create them one by one using the RetroRules webpage (https://retrorules.org/diy). As a result, we generated reaction rules for 5064 reactions with RetroPath RL (Figure 4a). Overall, PROXIMAL2 shows a higher level of automatization to generate reaction rules.

**Figure 4:**
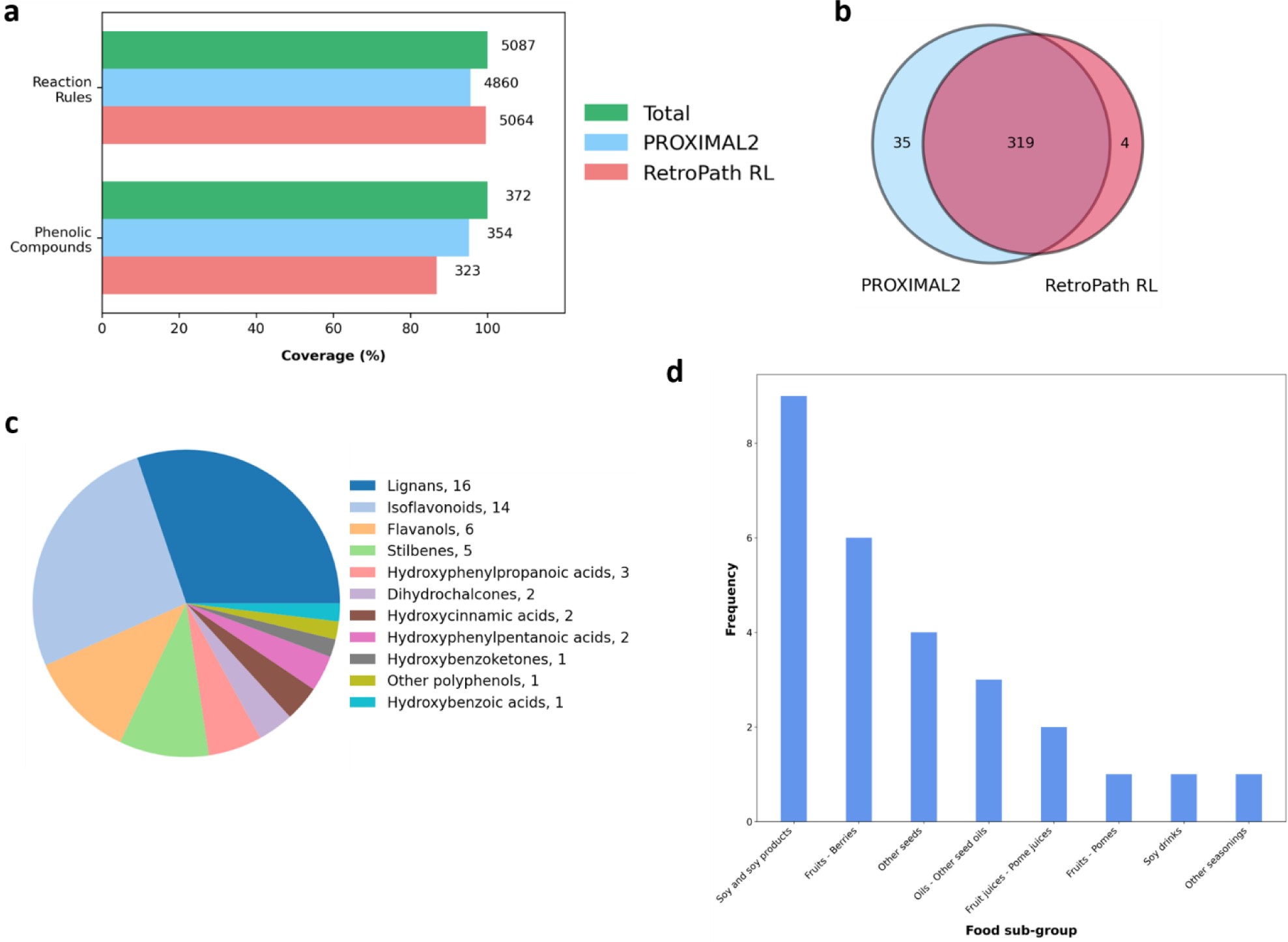
Comparison between PROXIMAL2 and RetroPath RL. **a)** Representation of reaction coverage regarding the ability to generate rules by the algorithms and the coverage of phenolic compounds to produce promiscuous products. The number to the right of the bars represents the absolute number of reactions and compounds; **b)** Venn diagram of the phenolic compounds that can be potentially degraded by PROXIMAL2 and RetroPath RL; **c)** Representation of the different sub-classes of the 53 phenolic compounds captured by PROXIMAL2. The number of compounds belonging to the sub-class is expressed in the legend, *e.g.* ‘Lignans, 16’. **d)** Frequency of sub-groups of foods associated with the 53 phenolic compounds exclusively predicted by PROXIMAL2 in comparison with (Balzerani *et al*., 2022).

We compared the reactions rules obtained with PROXIMAL2 and RetroPath RL, finding that 4837 were present in both cases, whereas 23 and 227 were unique to PROXIMAL2 and RetroPath RL, respectively (Figure 4a). The main differences between both approaches in the generation of rules are due to several reasons: (i) different treatment of cofactors (29.5%); (ii) ability of RetroPath RL to deal with stereochemistry (25.1%); (iii) reactions no longer present in the MetaNetX database, *i.e.* impossibility to download the reaction equation from MetaNetX and, consequentially, they were not considered with PROXIMAL2 (16.7%); iv) differences in the definition of MCS and reaction centres, *e.g.* the mandatory condition in the extraction of MCS in PROXIMAL2 (but not in RetroPath RL) that a match between atoms included in rings can happen only if the atoms are part of rings in both substrate and product (15%). As a result, RetroPath RL appears to cover a slightly wider area of the chemical space than PROXIMAL2.

Then, we applied the generated rules with both approaches to 372 phenolic compounds (Supplementary Table 4) obtained from Phenol-Explorer database (Rothwell *et al*., 2013). Since RetroPath RL provides all reaction products that can be generated from a query compound and PROXIMAL2 generates one product at a time, we filtered out the results for which PROXIMAL2 was not able to predict the whole set of putative products. We obtained results for 354 out of 372 phenolic compounds using PROXIMAL2, whereas RetroPath RL generated products for 323 (Figure 4a). Specifically, 319 of them were in common, while 35 were specific for PROXIMAL2 and 4 for RetroPath RL (Figure 4b). The differences observed between PROXIMAL2 and RetroPath RL are mainly caused by two factors: i) a different filter of chemical similarity for the query compound, namely PROXIMAL2 uses the Dice coefficient and RetroPath RL the Tanimoto coefficient; ii) differences in the definition of MCS, noted above, which determines a different reaction centre and local neighbourhood. These differences are emphasized by the fact that the overlap between PROXIMAL2 and RetroPath RL upon the generated products is only 60%. For this calculation, we assumed that both approaches reached the same product from phenolic compounds when the chemical similarity (Dice coefficient) was equal to 1.

In our previous work (Balzerani *et al*., 2022), we applied RetroPath RL to the same 372 phenolic compounds and extracted putative products for 303 of them using the recommended diameters by the authors (more than 6). Here, PROXIMAL2 could generate putative products for 53 additional phenolic compounds (301 were in common between PROXIMAL2 and RetroPath RL under this scenario). These 53 phenolic compounds were connected to 430 metabolites involved in AGREDA. Interestingly, we found that *Lignans* and *Isoflavonoids* were highly represented in this set of metabolites, namely 16 and 9 metabolites, respectively (Figure 4c). In addition, according to the food composition provided by Phenol-Explorer, the 53 phenolic compounds are part of 8 sub-groups of food, with *Soy and soy products* and *Fruits – Berries* the most annotated (Figure 4d). Overall, these results show that PROXIMAL2 complements our previous analysis with RetroPath RL and opens new research directions to understand the metabolism of phenolic compounds in the human gut microbiota.

## Discussion

Cellular metabolism involves the totality of chemical transformations that can occur in organisms and, though our knowledge is continuously growing, many metabolic pathways are still incomplete. A relevant case in the field of nutrition is the metabolism of phenolic compounds in the human gut microbiota, which remains largely unknown. Phenolic compounds, which are mainly derived from foods of plant origin (Scalbert, Johnson, *et al*., 2005; Scalbert, Manach, *et al*., 2005), are converted into bioactive metabolites that appear to limit the risk of several major diseases, such as coronary heart disease (Heim *et al*., 2002), cancer (Halliwell, 2002) or diabetes (Dembinska-Kiec *et al*., 2008). This fact has stimulated research to complete the knowledge about the degradation pathways of these nutrients in the human gut microbiota.

Recently, several methodologies have been developed to fill in metabolic gaps. The computational analysis of the metabolic space through enzyme promiscuity has received much attention. Specifically, rule-based methods have grown in number and quality in the last years (Ni *et al*., 2021). In a previous work (Balzerani *et al*., 2022), we applied a well-known rule-based enzyme promiscuity algorithm, RetroPath RL, to predict the degradation pathways of 372 phenolic compounds from Phenol-Explorer. Here, we explore a different rule-based algorithm, PROXIMAL, to address the same question. Given the current limitations and KEGG dependencies of PROXIMAL, we developed PROXIMAL2, which can automatically generate rules for a wider spectrum of reactions and make more reliable and comprehensive predictions for the degradation of phenolic compounds in the human gut microbiota.

As detailed in Methods section, without KEGG dependencies, PROXIMAL2 automatically extracts reactions rules without relying on the KEGG database, and replicates the look-up tables defined in PROXIMAL for predicting novel reactions. Further, PROXIMAL2 includes new features that were not part of PROXIMAL and expands its scope of application. In particular, PROXIMAL2 is able to capture complex transformations that involves multi step reactions through the development of multi-centres look-up tables. In addition, a detection of possible atom charges was implemented in PROXIMAL2, which allows tracking the charges present in substrates and products and adding or removing charges depending on the transformation. These new features of PROXIMAL2 significantly improved the performance of PROXIMAL, as described in the Results section with the comparison with KEGG reactions. PROXIMAL2 increased the coverage of KEGG reactions that can be potentially used for predicting enzyme promiscuity, namely 1950 KEGG reactions can be considered with PROXIMAL2 but not with PROXIMAL. Second, PROXIMAL2 shows higher accuracy than PROXIMAL in recovering annotated products from the substrates in KEGG reactions in the different scenarios considered in Figure 3a.

PROXIMAL2 was applied to predict novel degradation routes of phenolic compounds in the human gut microbiota. Note here that this study had not been feasible with PROXIMAL, given its KEGG dependences and the complexity of annotated reactions in AGREDA for phenolic compounds. We compared the results of PROXIMAL2 with RetroPath RL. With respect to rule generation, we found that PROXIMAL2 is more practical in the automatic extraction of reaction rules, as illustrated with the subset of manually curated reactions included in the metabolic space. On the other hand, RetroPath RL covers a wider range of chemical space, obtaining reaction rules for 227 reactions for which PROXIMAL2 could not obtain look-up tables. Although the difference is limited, we expect to address these limitations in future developments.

In addition, we found that PROXIMAL2 and RetroPath RL have a significant overlap in the subset of phenolic compounds for which they could find a putative product. Both algorithms obtained a promiscuous product in 319 out of 372 compounds (85.7%). However, we found PROXIMAL2 and RetroPath RL complementary; PROXIMAL2 predicted new output products for 35 phenolic compounds that were not captured by RetroPath RL. We found that these differences are dependent on the choice of specific parameters, *e.g.* diameter in RetroPath RL or chemical similarity coefficient used in both approaches, which emphasizes the importance of investigating the optimal set of parameters for different algorithms. Importantly, our results open new research directions in the metabolism of phenolic compounds in the human gut microbiota. We expect to further analyse and experimentally validate the results obtained from this study.

Finally, beyond the problem of phenolic compound degradation in the human gut microbiota, PROXIMAL2 is a general-purpose algorithm and can be applied to predict novel associations of other molecules of interest to putative products. PROXIMAL2 can be complementarily used with other existing rule-based methods, such as RetroPath RL, to provide novel insights into metabolic gaps for different applications.

## Data Availability

The data employed in this study can be obtained from the following databases: (i) Metabolic model: AGREDA v1.1.0 (https://github.com/francesco-balzerani/AGREDA_1.1); (ii) Metabolites and Chemical rules: PubChem (https://pubchem.ncbi.nlm.nih.gov/), Human Metabolome Database (https://hmdb.ca/), KEGG (https://www.genome.jp/kegg/), MetaNetX (https://www.metanetx.org/), RetroRules (https://retrorules.org/), Phenol-Explorer (http://phenol-explorer.eu/). The source data underlying Figs. 3 and 4 are provided as Supplementary Tables 1, 2, 3, 4.

## Code Availability

Python code of PROXIMAL2 is available in https://github.com/HassounLab/PROXIMAL2

## Author contributions

F.J.P. and S.H. conceived this study. F.B., T.B., S.P-B., L.V., F.J.P., and S.H. developed the algorithm and performed the computational analysis. All authors wrote, read, and approved the manuscript.

