## Supplementary Information for "Extending PROXIMAL to predict degradation pathways of phenolic compounds in the human gut microbiota"

### Supplementary Methods: PROXIMAL2: a new enzyme promiscuity method for complex multi-step reactions

**Selected parameters in the Maximum Common Substructure algorithm in RDKit**

In the step 3 of the PROXIMAL2 pipeline, detailed in the main text, substrate-product pairs are aligned with the maximum common substructure (MCS) algorithm that is available in RDKit (Landrum, 2011): function *findMCS* in the module *rdFMCS*. We made use of 3 specific parameters:

- *bondCompare*. It is defined as *CompareOrderExact*, which ensures that the common substructures share the same bond type and order;

- *ringMatchesRingOnly*. It is defined as *True* and it guarantees that atoms belonging to rings in substrates match only with atoms belonging to rings in products. Supplementary Figure 1a shows an example MCS where this parameter is not considered, leading to a meaningless match between atoms in a linear structure in the substrate, C00353, and atoms belonging to a ring in the product, C20158. This issue is corrected in Supplementary Figure 1b, where this parameter is introduced. Moreover, the parameter provides the capacity of the algorithm to extract the partial match between rings, as shown in Supplementary Figure 1c, where the alignment between atoms of the rings is not complete;

- *matchValence*. This parameter is set as *True* and it considers the valence of the atoms in the MCS, discarding a match between atoms with a different valence. Supplementary Figure 1d shows a case when the difference in valence of the atoms determines the partial match of the substructures. In particular, the atom 1 in the substrate, C00019, and atom 14 in the product, C00021, are the same atom, but they differ in valence and, therefore, the correspondence is denied.

**a**

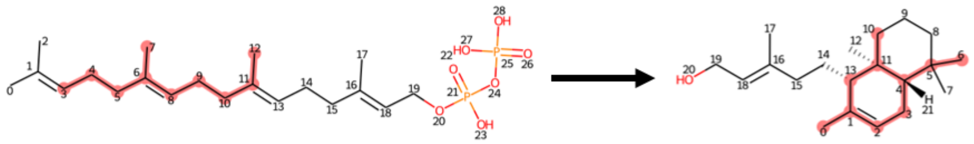

C00353 C20158

| Alignment via MCS | C00353 | 3 | 4 | 5 | 6 | 7 | 8 | 9 | 10 | 11 | 12 |
|---|---|---|---|---|---|---|---|---|---|---|---|
| | C20158 | 10 | 11 | 13 | 1 | 0 | 2 | 3 | 4 | 5 | 6 |

**b**

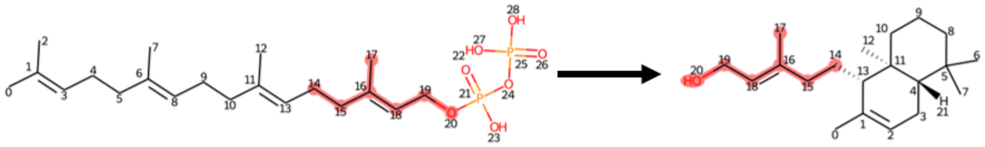

C00353 C20158

| Alignment via MCS | C00353 | 14 | 15 | 16 | 17 | 18 | 19 | 20 |
|---|---|---|---|---|---|---|---|---|
| | C20158 | 14 | 15 | 16 | 17 | 18 | 19 | 20 |

**c**

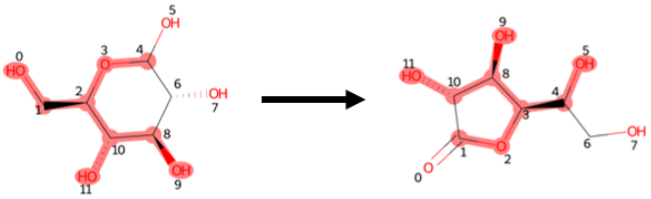

C00031 C03107

| Alignment via MCS | C00031 | 0 | 1 | 2 | 3 | 4 | 8 | 9 | 10 | 11 |
|---|---|---|---|---|---|---|---|---|---|---|
| | C03107 | 5 | 4 | 3 | 2 | 1 | 10 | 11 | 8 | 9 |

**d**

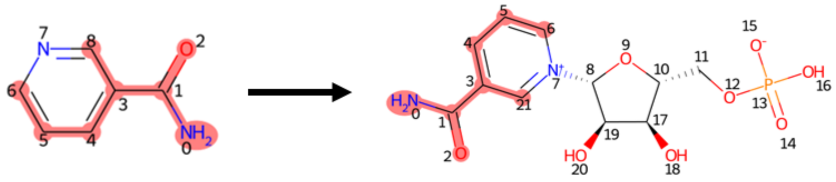

C00153 C00455

| Alignment via MCS | C00153 | 0 | 1 | 2 | 3 | 4 | 5 | 6 | 8 |
|---|---|---|---|---|---|---|---|---|---|
| | C00455 | 0 | 1 | 2 | 3 | 4 | 5 | 6 | 21 |

**Supplementary Figure 1:** Illustration of different parameters in the maximum common substructure (MCS) algorithm. **a)** Resulting MCS for the KEGG reaction *R09874* with the parameter *ringMatchesRingOnly* inactive; **b)** Resulting MCS for the KEGG reaction *R09874* with the parameter *ringMatchesRingOnly* active; **c)** Representation of the partial match between subgraph of the rings in the substrate and product due to the parameter *ringMatchesRingOnly*. KEGG reaction id: *R00301*); **d)** Resulting MCS with the parameter *matchValence* active, which discards matching between atoms with a different valence. KEGG reaction: *R00828*. In all the cases MCS is shown in red colour.

**Bond rearrangement between atoms belonging to the Maximum Common Substructure**

In multi-step reactions the substrate goes through several structural transformation, as reflected in Step 4 in the PROXIMAL2 pipeline. In some cases, structural changes can involve bonds between atoms belonging to the Maximum Common Substructure, which is not reflected in standard 'key' tables. Changes include the order of a bond (Supplementary Figure 2a), and a removal/formation of a bond (Supplementary Figure 2b). In order to consider these changes in the look-up tables, the bonds between atoms included in the MCS are analysed and compared between the substrate and the product, checking for differences and modifications. Then, similarly to reaction centres, the atoms involved in the bonds and their respective adjacent and distant neighbours are extracted.

**a**

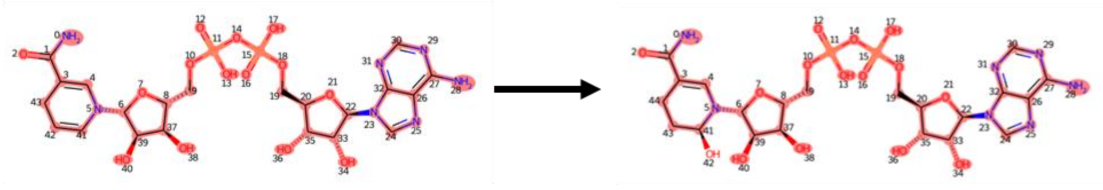

C00004 C04856

| Alignment via MCS | C00004 | 0 | 1 | 2 | 3 | … | 40 | 41 | 42 | 43 |
|---|---|---|---|---|---|---|---|---|---|---|
| | C04856 | 0 | 1 | 2 | 3 | … | 40 | 41 | 43 | 44 |

**b**

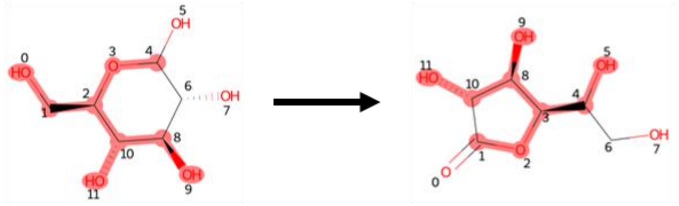

C00031 C03107

| Alignment via MCS | C00031 | 0 | 1 | 2 | 3 | 4 | 8 | 9 | 10 | 11 |
|---|---|---|---|---|---|---|---|---|---|---|
| | C03107 | 5 | 4 | 3 | 2 | 1 | 10 | 11 | 8 | 9 |

**Supplementary Figure 2:** Representation of bond changes between atoms belonging to the MCS. **a)** Definition of the MCS for the KEGG reaction *R00129.* The bond between atom 41 and 42 in the substrate (C00004) is a double bond and the corresponding bond between atom 41 and 43 within the product (C04856) is a single bond due to the addition of the functional group. KEGG reaction: *R00129*. The MCS is not completely included. However, from atom 0 to atom 41 the alignment is numerically exact in the two molecules; **b)** Resulting MCS for the KEGG reaction *R00301*. The bond between atom 10 and 1 in the product (*C03107*) is not present on the substrate (C00031), whose corresponding atoms are number 4 and 8. This is a multi-step reaction, where, in addition to other structural changes, it can be observed the hexagon ring cleavage and the formation of the pentagon ring within the MCS.
